# Emotional intensity can enrich or degrade memories: impact of the amygdalar pathway on hippocampus through inhibitory neurons

**DOI:** 10.1101/2022.03.17.484812

**Authors:** Yohan J. John, Jingyi Wang, Daniel Bullock, Helen Barbas

## Abstract

The brain’s emotional system powerfully modulates processing of context and episodic memory. A key pathway that mediates these effects is the projection from the amygdala to the hippocampus. Wang and Barbas (1) uncovered a distinctive pattern in the pathways from amygdala to hippocampus in primates. In hippocampal CA3, a pathway from the amygdala innervated excitatory pyramidal neurons as well as parvalbumin (PV) and calretinin (CR) inhibitory neurons. In hippocampal CA1, amygdalar projections also innervated pyramidal neurons and CR interneurons, but not PV interneurons. The effects of these complex circuits can best be probed using computational simulations. We developed a model of spiking neurons to investigate the implications and significance of these amygdala-hippocampal circuits for affective influence on processing mnemonic context, and to test their effects as input from the amygdala gradually increased. Our simulations revealed that moderate input from the amygdala can enhance detail in CA3 representations that can correctly sort out contexts and episodes from memory. However, high amygdalar input suppressed CA3 responses to non-amygdalar inputs through powerful inhibitory neurons, leading to memory representations that lack detail. Moreover, high amygdalar input prematurely hastened the timing of responses in CA1 occurring when the current situation broadly and non-specifically matched a remembered context. Amygdalar pathways to hippocampus enable a mechanism whereby affective signaling appropriately enhances hippocampal representations of remembered context. However, when amygdalar input is excessive in high emotional arousal, there is loss of memory detail and overgeneralization, as seen in post-traumatic stress disorder or pathologic phobias.

## Introduction

Emotion and memory are deeply intertwined. It is well known that emotionally salient events and episodes can leave deep traces in memory (2). However, abnormally intense emotional experiences, such as traumas, may instead weaken memory of episodic details, or contribute to overgeneralization in neutral situations. The right amount of emotion can render memories vivid and long-lasting, but excessive emotion can degrade other types of information. A key question, then, is how the neural circuit that brings emotion and memory together can lead to this spectrum of effects, ranging from memory enhancement to loss of episodic detail and specificity.

The amygdala and hippocampus are central nodes in this affective-memory circuit. The amygdala is a well-established hub of social and affective processing (3–7) and can have powerful effects on perception and cognition via its potent projections to thalamus and cortex (8– 12). The hippocampus, and in particular the anterior hippocampus (ventral in rodents), is crucial to the formation of memories of episodes and contexts (13–15). A seminal line of research implicating the hippocampus in memory began with the amnesic patient known during his lifetime as ‘HM’. The severe anterograde amnesia that HM, Henry Molaison, displayed was attributed to the removal of both his hippocampi as part of a treatment for severe and drug unresponsive epilepsy. Postmortem analysis showed that some parts of hippocampus had been spared, but the basolateral amygdala of both hemispheres had been completely ablated (16). This suggested a structural basis for a symptom that was sometimes neglected in popular accounts: HM had almost no fear, and also displayed a muted ability to experience hunger and thirst (17). Additional human case studies of more localized lesions have confirmed this functional division of labor between amygdala and hippocampus (18).

Decades of research have fleshed out this understanding of the amygdala and hippocampus, but their interactions remain incompletely characterized. To this end, Wang and Barbas investigated the nature of the projections from the amygdala to hippocampus in rhesus monkeys (1).

Projections from the basolateral amygdala terminated across the entire hippocampus, but most were found in the anterior hippocampus, and predominantly in the upper layers of the CA3 and CA1 subfields. The pattern of communication between the amygdalar pathway to these hippocampal subfields suggested a mechanism through which the amygdala could boost hippocampal memory, but also reduce episodic and contextual detail during high emotional arousal.

Specifically, in CA3 the amygdalar pathway innervated pyramidal neurons as well as inhibitory interneurons labeled with the calcium-binding proteins calretinin (CR) or parvalbumin (PV). In CA1, the amygdala also projected to pyramidal and CR neurons, but not to PV neurons. The amygdalar projection to hippocampus was very strong, as revealed by density, synaptic specializations and significantly larger terminals than seen in nearby unlabeled synapses that did not receive direct projections from the amygdala. Some implications of this connectivity pattern were suggested in a conceptual model (1). Here we tested the implications of this complex circuit with the help of a computational model of the amygdala-hippocampus circuit. Our spiking circuit model revealed how emotional salience signals sent from the amygdala can regulate the level of mnemonic detail represented in CA3. These signals also modulate the ability of CA1 to compare mnemonic patterns in CA3 with input from entorhinal cortex (EC) that conveys information about the current context of the organism.

The spiking model implements the basic circuit motif described above, supplementing it with information about the local connectivity within CA3 and CA1 derived from the broader literature. We posited that in both CA3 and CA1, the PV neurons serve as sources of strong perisomatic inhibition of nearby pyramidal neurons, and that the CR neurons primarily disinhibit the same pyramidal neurons: they suppress interneurons that inhibit the dendrites of pyramidal neurons. This role of CR interneurons has been observed in the upper layers of cortex, and evidence suggests it holds for the hippocampus as well (19,20). In both CA3 and CA1, CR neurons are assumed to mediate a broad pattern of disinhibition. We also assumed that the PV interneurons mediate lateral inhibition of nearby pyramidal neurons in CA3, as described (21,20,22).

Our simulations showed that moderate activation from the amygdala can disinhibit CA3 pyramidal neurons, providing CA1 with rich ‘recollections’ that can be compared with the current situation. When amygdala activity is excessive, disinhibition via CR neurons of CA3 pyramidal neurons is overwhelmed by lateral inhibition acting via the powerful PV neurons. As a consequence, high amygdala activity, which is assumed to accompany abnormally intense emotional experiences, can cause loss of detail in CA3 representations of episodic memories, and disrupt the role of CA1 as a comparator of current context and memories.

## Methods

### Model design

The basic connectivity of the computational model is based on the conceptual circuit derived from detailed data on the amygdala-hippocampus pathway (1). Amygdalar synapses in hippocampus were found to be strong in terms of density and terminal size. Moreover, the amygdalar pathway had specializations of double synapses onto the same dendritic segment – a rare occurrence anywhere in the brain (Figure 1). The basic amygdalar connectivity in hippocampus is shown schematically in Figure 2. The neurons in CA3 are divided into 5 ensembles of which 2 are shown in Figure 2, for clarity. Each ensemble consists of excitatory pyramidal neurons, inhibitory PV interneurons, inhibitory CR interneurons, and dendrite-inhibiting (DI) interneurons. PV interneurons inhibit the cell bodies of pyramidal neurons within their own ensembles, as well as pyramidal neurons in neighboring ensembles, enabling both self-inhibition and lateral inhibition at the level of pyramidal neuron ensembles. Each PV interneuron also inhibits all other PV interneurons. DI interneurons project only within their ensembles, inhibiting pyramidal neurons. The CR interneurons inhibit DI interneurons within their own ensembles as well as in neighboring ensembles, enabling widespread disinhibition of CA3 pyramidal neurons. One of the ensembles receives input from the amygdala. In this ensemble, the pyramidal neurons, CR, and PV interneurons receive excitatory input from the amygdala.

**Figure 1.**
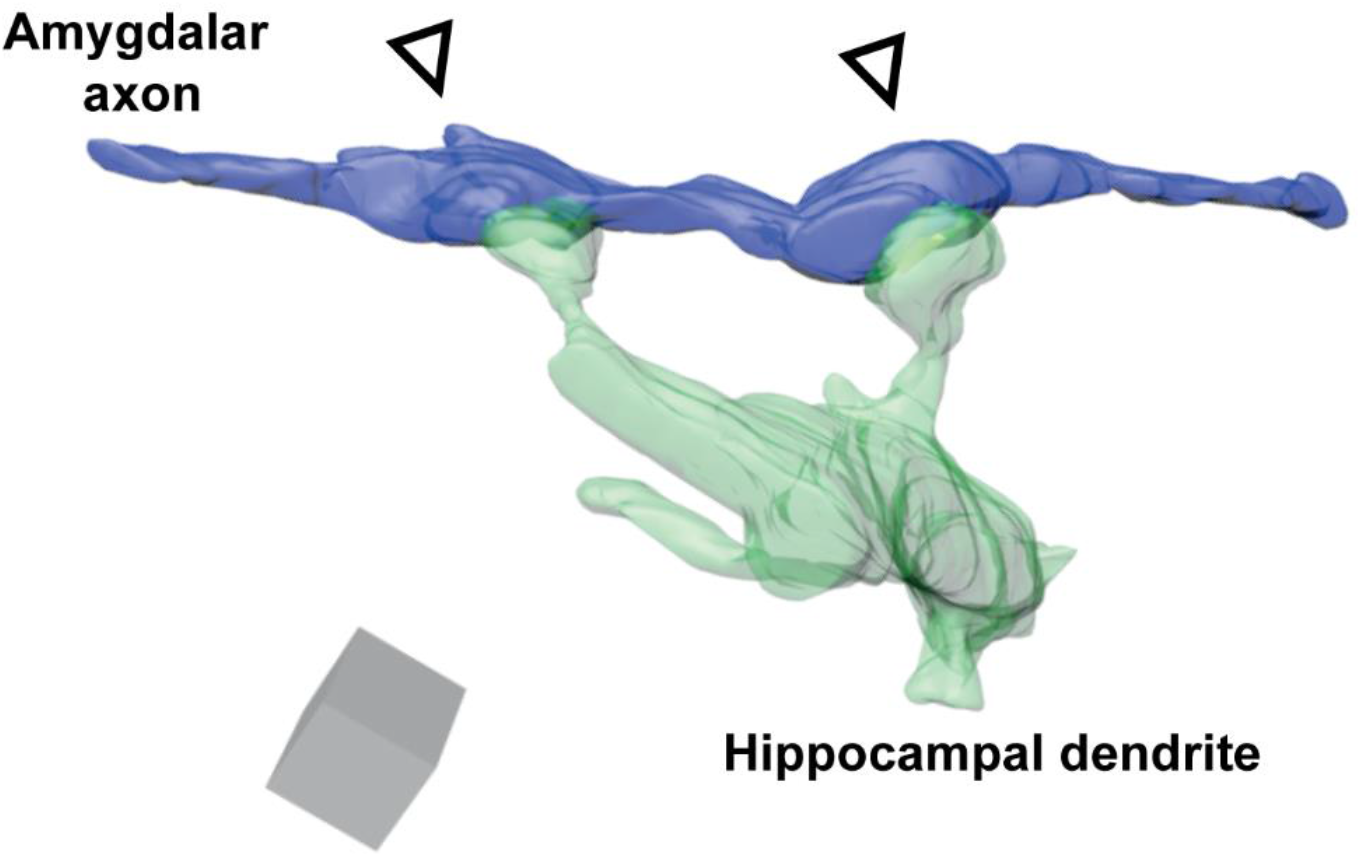
Example of the strength of the pathway from the amygdala to hippocampus. The figure shows a reconstruction of images captured with an electron microscope from serial uninterrupted sections (50 nm each) through the synapse. Two boutons on the same amygdalar axon (blue) form synapses with spines from the same dendritic segment (green) of a postsynaptic pyramidal hippocampal neuron in CA3. Such double synapses, which are rare anywhere in the brain, were frequently observed in the amygdalar pathway to hippocampus. Grey box shows scale: 1×1×1 micron. 3D reconstruction of images through the synapse was adapted from (1).

**Figure 2.**
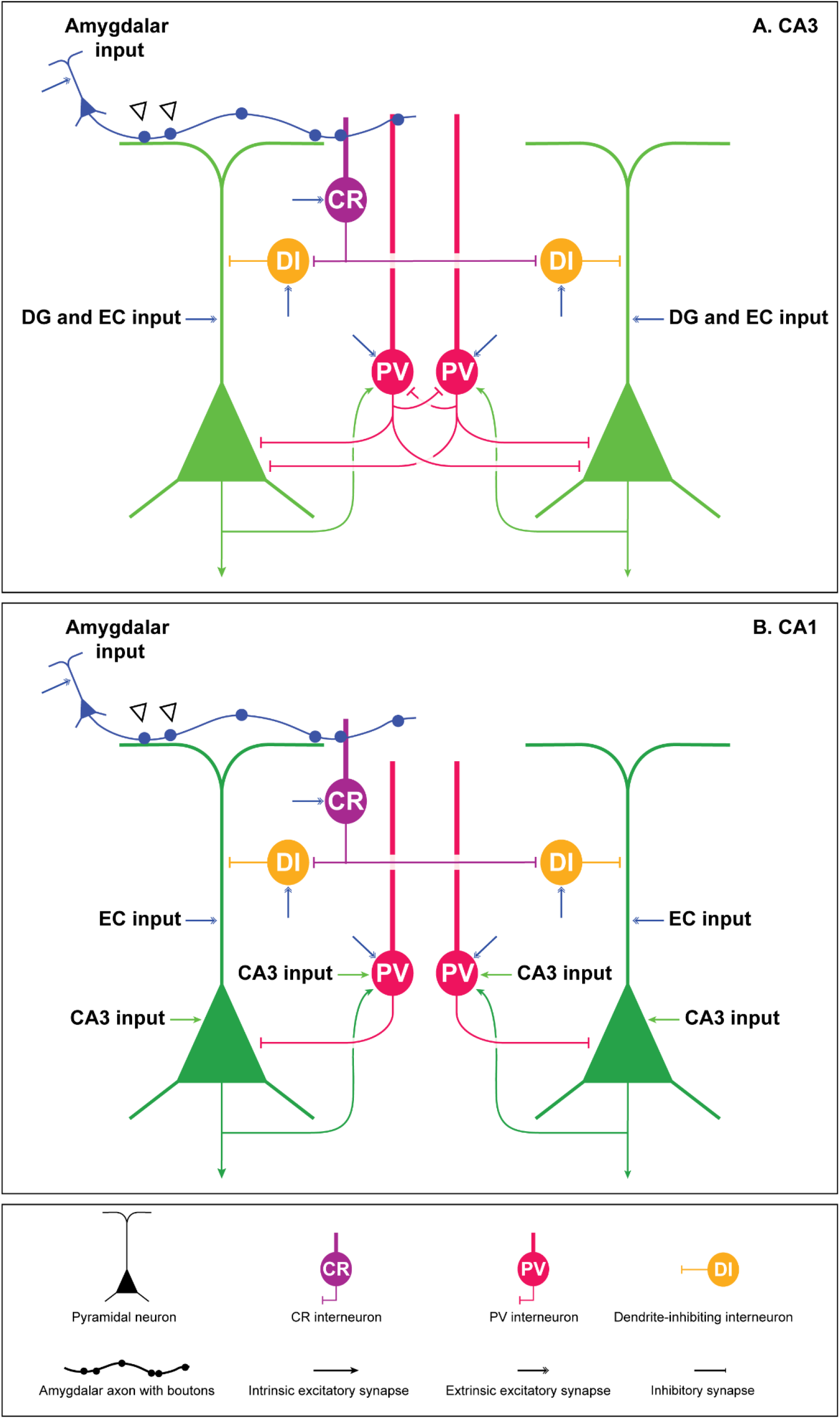
Schematic diagram of model circuit. A: The CA3 circuit contains five ensembles of neurons, two of which are shown here for simplicity. Each ensemble contains a population of pyramidal neurons, PV interneurons, CR interneurons, and dendrite-inhibiting (DI) interneurons. Only one of the five ensembles receives amygdalar input, which arrives at the pyramidal neurons, PV, and CR interneurons. B: The CA1 circuit consists of two ensembles of neurons. Each ensemble contains a population of pyramidal neurons, PV interneurons, CR, interneurons, and dendrite-inhibiting (DI) interneurons. Amygdalar input arrives at one of the two ensembles. CA3 pyramidal neurons send feedforward excitation to CA1 pyramidal neurons and PV interneurons. This wiring pattern is based on information from the literature as well as our findings on the pathways from the amygdala to hippocampus in rhesus monkeys (1).

The impact of the level of the amygdalar input on CA3 is shown schematically in Figure 3.

**Figure 3.**
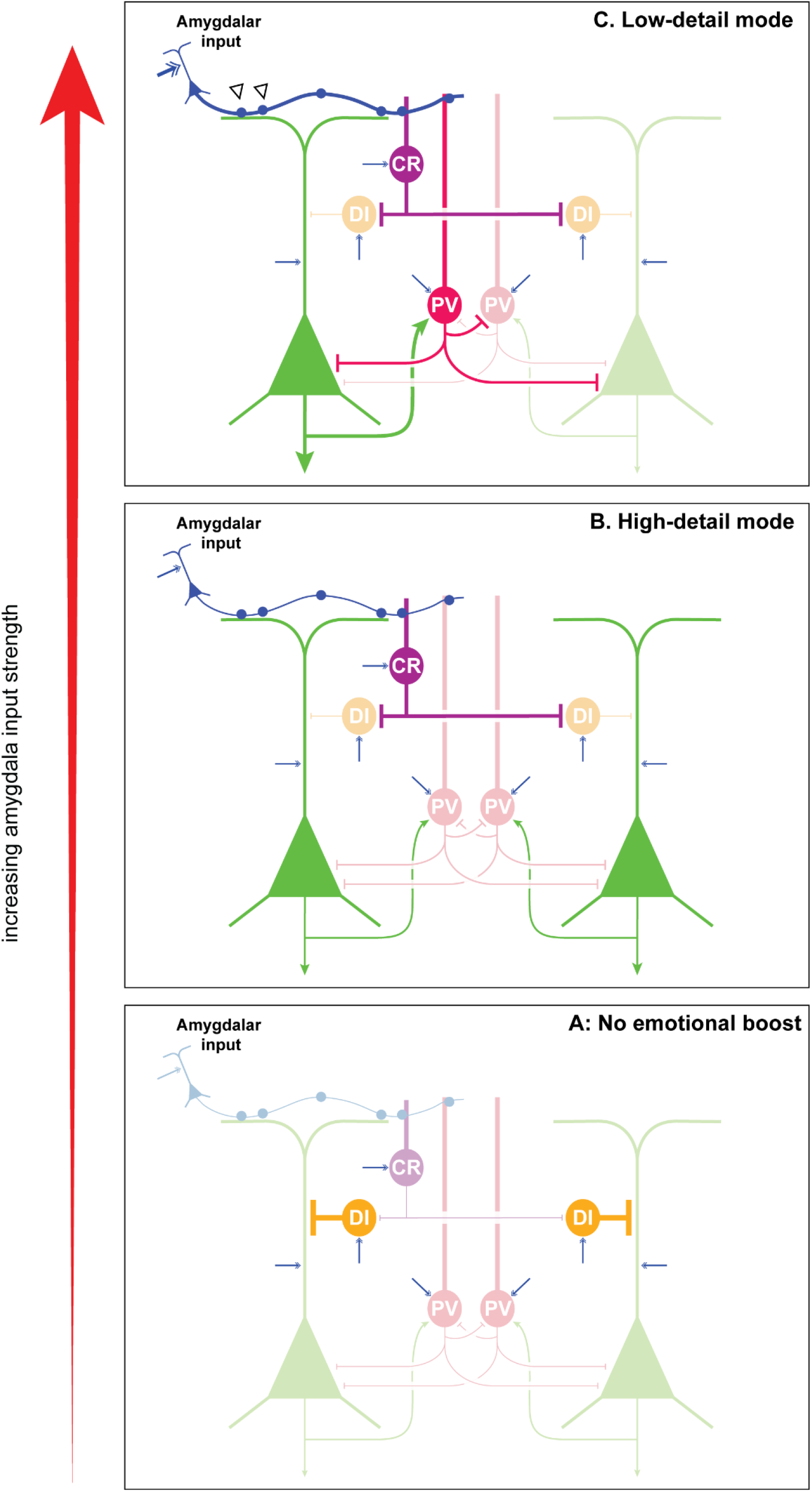
Schematic diagram of amygdalar modulation of information detail in CA3. A: For low levels of amygdalar input, dendrite-inhibiting (DI) interneurons are not inhibited by CR interneurons, leading to low firing in CA3. B: For moderate levels of amygdalar input, DI interneurons are inhibited by CR interneurons, causing disinhibition of CA3 pyramidal neurons, which corresponds to a high detail state triggered by an emotional boost. C: For abnormally high levels of amygdalar input, PV interneurons begin to fire very strongly, leading to strong inhibition of pyramidal neurons (light green). This leads to a low detail state. Only pyramidal neurons receiving amygdalar input withstand this strong inhibition (dark green).

The CA1 subfield is divided into two ensembles, one of which receives amygdalar input. The local circuitry in CA1 is analogous to that of CA3. CA1 pyramidal neurons receive dendritic excitation from amygdala and EC, and somatic excitation from CA3. The CA1 CR interneurons in one of the ensembles receive excitation from the amygdala. Each CA1 pyramidal neuron receives excitation from all CA3 pyramidal neurons. CA1 PV interneurons also receive excitation from all CA3 pyramidal neurons (23). All CA3 and CA1 neurons receive tonic extrinsic excitatory input in the form of Poisson spikes. “Extrinsic” here implies that these sources of excitation are not explicitly modeled. Intrinsic inputs arise via the circuit connections depicted in Figure 2.

An important component of the CA1 pyramidal neurons is the M-current, which mediates a switch between tonic and phasic modes of firing. We base this mechanism on a recent detailed biophysical model (24). The phasic mode occurs when a CA1 pyramidal neuron is ‘primed’ by excitation in its apical dendritic compartment, which opens KCNQ channels while preventing firing. When a subsequent driving input arrives at the somatic compartment, the neuron releases a phasic volley of spikes and then becomes silent as a result of the hyperpolarizing effects of the M-current. If the driving input arrives without prior priming, then the neuron shows tonic firing. The phasic mode signals a match between the priming input (from EC and amygdala) and the driving input (from CA3). The tonic mode signals a mismatch between the priming and the driving inputs.

Driving input serves as a form of prior expectation, while the priming input conveys information about the current situation of the organism. If we interpret input from EC as signaling the present context of events, and input from CA3 as signaling memories that have been elicited, then a match between EC and CA3 indicates that the current situation is familiar, as it resembles prior episodic or contextual memories. The phasic match signal can trigger responses that were previously associated with the remembered situation. A mismatch between EC and CA3 indicates that none of the memories is a good enough template for the current context. The tonic mismatch firing can trigger a variety of responses, including alerting, heightened attention, learning of new episodic memories, and/or disengagement from ongoing behaviors. The CA1 match/mismatch mechanism is depicted schematically in Figure 4.

**Figure 4.**
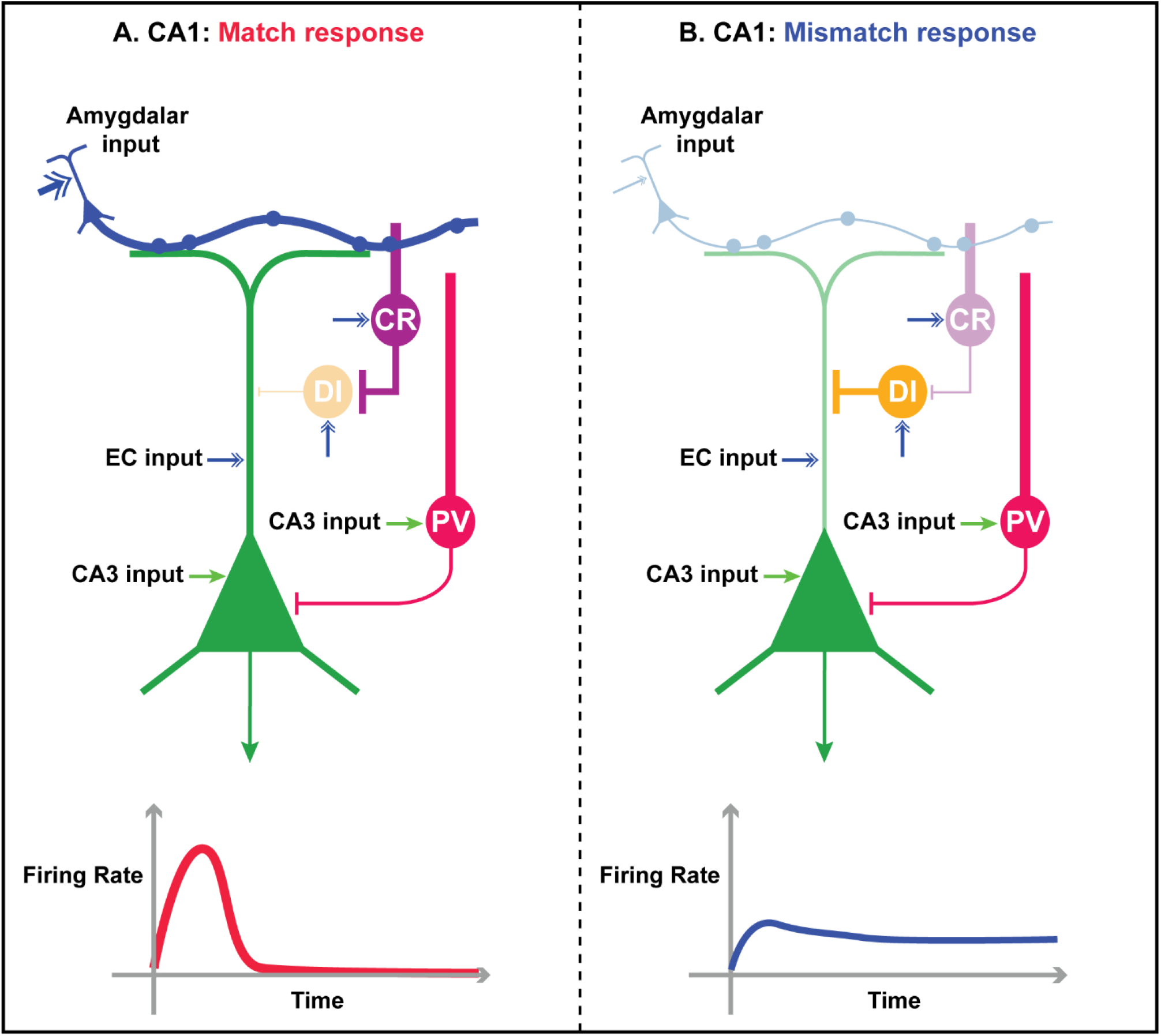
Schematic diagram of match/mismatch mechanism in CA1. A: If priming input from the amygdala and EC is present prior to driving input from CA3 pyramidal neurons, the CA1 pyramidal neuron produces a phasic response, corresponding to a match (bottom, red trace). B: If there is insufficient priming from the amygdala, the CA1 pyramidal neuron responds with tonic firing, corresponding to a mismatch (bottom, blue trace).

### Simulation details

Neurons were modeled using the Izhikevich spiking framework (25,26) in combination with realistic AMPA, NMDA and GABA synapses. Each neuron produced a fast synaptic effect (AMPA for excitatory neurons, and GABAA for inhibitory neurons) and a slow synaptic effect (NMDA for excitatory neurons and GABAB for inhibitory neurons). This modeling framework has been described in detail in prior studies (11,27). Pyramidal neurons and DI and CR interneurons were modeled as regular-spiking neurons, which have type-I firing rate responses, i.e., they do not have a sharp firing threshold. PV interneurons were modeled as fast-spiking neurons, which have type-II firing responses, i.e., they have a threshold below which input cannot trigger spiking. Each pyramidal neuron in CA3 and CA1 was modeled as having two compartments. Pyramidal neurons in CA1 possessed KCNQ channels, predominantly in the somatic compartment. Extrinsic excitatory inputs to the neurons were modeled as Poisson spike trains that produce the same slow and fast synaptic effects as mentioned above. Simulations were performed in MATLAB (R2021b). A complete mathematical description of the model and the extrinsic inputs is provided in the Supplementary Material.

## Results

The population of CA3 neurons was divided into 5 ensembles, one of which received amygdalar input. Similarly, the CA1 was divided into two ensembles, one of which received amygdalar input. Each ensemble contained pyramidal neurons, dendrite-inhibiting interneurons, soma-inhibiting PV interneurons, and disinhibitory CR interneurons, which inhibit the dendrites of inhibitory neurons that innervate adjacent excitatory (pyramidal) neurons. The simulations of the integrated amygdala-CA3-CA1 model show how amygdalar inputs affect both the level of detail in CA3 and the comparison process mediated by CA1. As the spiking rate of amygdala neurons was increased, the pyramidal neurons in CA3 were initially disinhibited via CR interneurons.

Further increases in amygdalar input recruited strong perisomatic inhibition from PV interneurons, causing inhibition to dominate, as depicted schematically in Figure 3. For the strongest amygdalar input levels, only the activity of pyramidal neurons receiving amygdalar input “survived”. This constitutes a low detail state in CA3, because of the silencing of the comparatively weaker input to CA3 from other brain sources. The downstream comparison process in CA1, in which the CA3 representation is compared with the current information from EC, is depicted schematically in Figure 4.

Figure 5 shows the spike rasters of the simulated CA3 and CA1 pyramidal neuron cell bodies from an example trial. Each trial consisted of a set of 5 simulations (A - E) that vary in the strength of amygdalar input. For all simulations within a trial, the neuronal parameters and input trains were identical, other than the strength of input from the amygdala. The pink horizontal bar in each subplot indicates the CA3 pyramidal neuron ensemble that receives direct amygdalar excitation. Moving from the bottom subplot (Figure 5A) to the top subplot (Figure 5E), the spiking rate of the amygdalar input was increased. In Figure 5A, the pyramidal neurons had a low response in CA3. This is because the inhibition of CA3 pyramidal neurons by dendrite-inhibiting interneurons was dominant, and the amygdalar input was insufficient to cause disinhibition via CR interneurons. As the amygdalar input was increased, the CA3 dendrites of pyramidal neurons became disinhibited, enabling all CA3 pyramidal ensembles to become maximally active (Figure 5C). This corresponds to the optimal detailed state of CA3, depicting a boost for affective significance from the amygdala, as well as event-related information from other brain sources. Further increases in amygdalar input (Figure 5D, E) caused inhibition via PV interneurons in CA3 to dominate, so that the CA3 pyramidal neurons that did not receive direct amygdalar input became comparatively suppressed. At the highest amygdalar input strength simulated (Figure 5E), only the ensemble that received direct amygdalar input remained strongly active in CA3 (highlighted with a pink bar). This was the state of CA3 that conveyed the least detail to CA1. The onset times of inputs to key neural populations were staggered in the simulations (indicated in Figure 5 with colored vertical bars). EC inputs to CA1 arrived first, followed by input to CA3, followed by amygdalar inputs to CA3 and CA1. The reasoning behind this temporal sequence is provided in the Supplementary Material.

**Figure 5.**
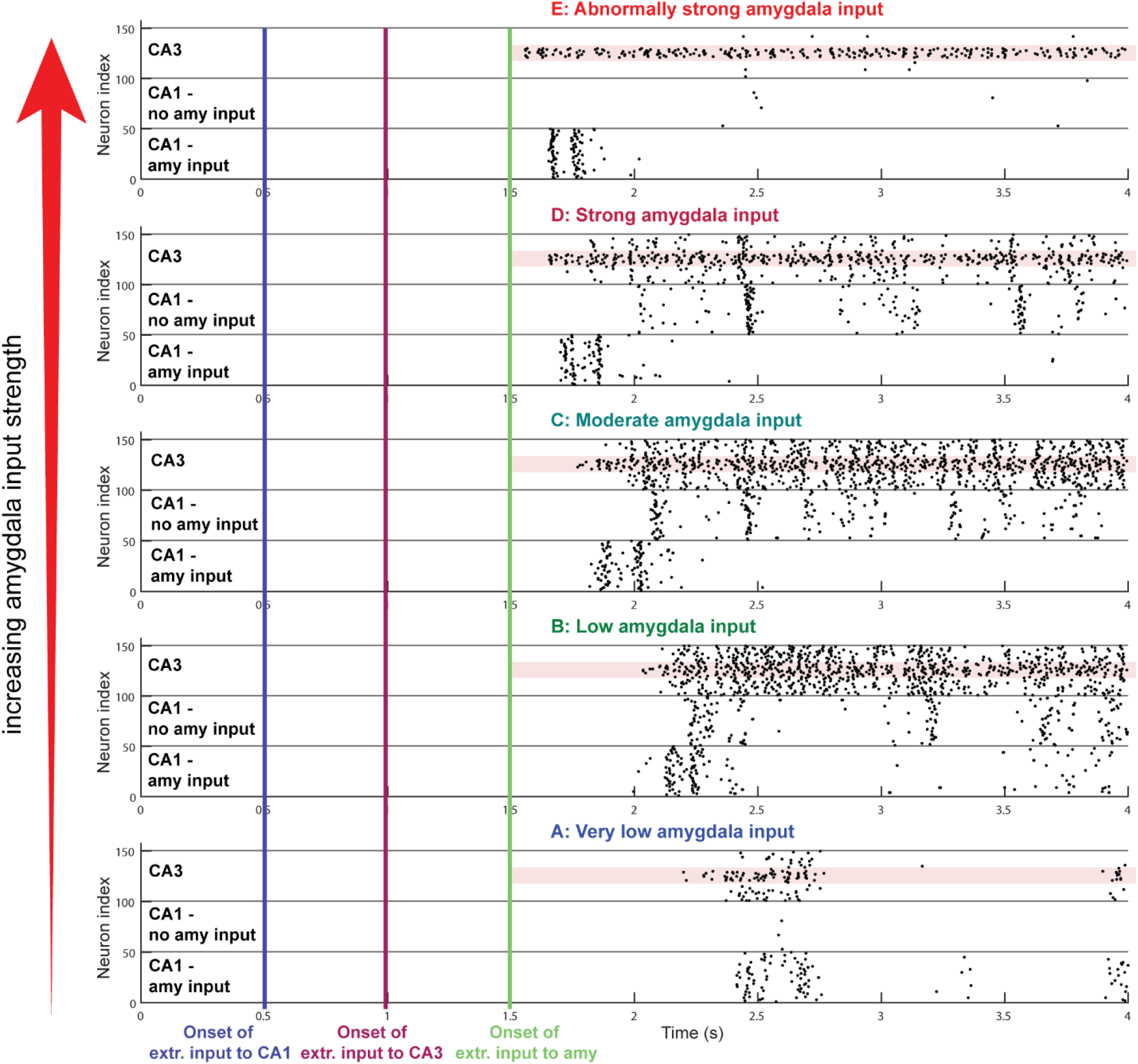
Simulation results: raster plot of spiking in CA3 and CA1 pyramidal neurons. Proceeding from the bottom plot to the top (A-E), the strength of input to hippocampus from the amygdala increases. Pink horizontal stripes indicate the CA3 pyramidal neuron subpopulation that receives amygdalar excitation. Vertical colored bars indicate onsets of extrinsic Poisson input to CA1 (blue), CA3 (purple), and amygdala (green). A: When there is low input from the amygdala, CA3 activity is weak and intermittent, as inhibition from dendrite-inhibiting interneurons is dominant. CA1 activity in the subpopulation that receives amygdalar input is also weak and intermittent, and in the subpopulation that receives no amygdalar input, activity is even weaker. B: As amygdalar input increases, CA3 CR interneurons inhibit the dendrite-inhibiting interneurons, disinhibiting the CA3 pyramidal neurons. CA1 pyramidal neurons begin to display the qualitative distinction between phasic responses (amygdalar input subpopulation) and tonic responses (no amygdalar input subpopulation). C: For moderate amygdalar input, CA3 pyramidal neurons are maximally disinhibited, leading to a high detail state. CA1 shows a clear qualitative distinction between the phasic match response (amygdalar input subpopulation) and the tonic mismatch response (no amygdalar input subpopulation). D: As amygdalar input increases further, inhibition from PV interneurons in CA3 becomes strong. The CA3 pyramidal neurons that receive amygdalar input (pink bars) are able to withstand the PV inhibition and remain strongly active. CA1 pyramidal neurons show the same qualitative distinction as in the moderate input case (C). E: When amygdalar input is abnormally strong, only the CA3 pyramidal neurons that receive direct amygdalar input “survive” the strong PV neuron inhibition (pink stripe). The other CA3 pyramidal subpopulations display very low firing rates (above and below pink stripes). CA1 pyramidal neurons that receive amygdalar input continue to show a phasic response, whereas the subpopulation that does not receive amygdalar input is suppressed as a result of inhibition as well as input from CA3 that is weaker than normal.

Figure 5 also shows how increasing amygdalar input affects firing patterns in CA1. The CA1 pyramidal neurons, which receive input from CA3, were divided into two subpopulations: both subpopulations received EC priming, but only one received amygdalar priming. This allowed us to highlight the possible consequences of the amygdala-CA3 effect downstream. Priming signals excited the apical dendrites of CA1 pyramidal neurons. Under normal circumstances this dendritic priming is too weak to elicit appreciable firing unless there is coincident driving input from CA3 pyramidal neurons, which arrives at the cell body. The CA1 pyramidal neurons displayed two qualitatively different modes that depended on the strength of priming to the apical dendrite (shown schematically in Figure 4). If a CA1 pyramidal neuron received sufficient priming, it responded to strong driving input from CA3 with a phasic volley of spikes followed by near-quiescence (Figure 5: lower CA1 ensemble in each subplot B-D).

Figure 6 summarizes the effect of increasing amygdalar input on CA3 activity, and also shows that the effects are robust with respect to small changes in initial conditions and Poisson input trains to each neuron. Data were averaged from 20 trials, each analogous to the trial shown in Figure 5. For each trial, each neuronal parameter and connection weight was drawn from a uniform distribution with a width of ±2.5% around the mean value shown in the table of parameters (Supplementary Table 1). The plots show that moderate amygdalar input allowed for appreciable activity in all the CA3 pyramidal neurons. As the strength of amygdalar input was increased, the activity in CA3 pyramidal neurons became more and more biased: for the highest level of amygdalar input simulated, the pyramidal neurons that did not receive direct amygdala excitation were suppressed, resulting in an impoverished, low-detail representation in CA3.

**Figure 6.**
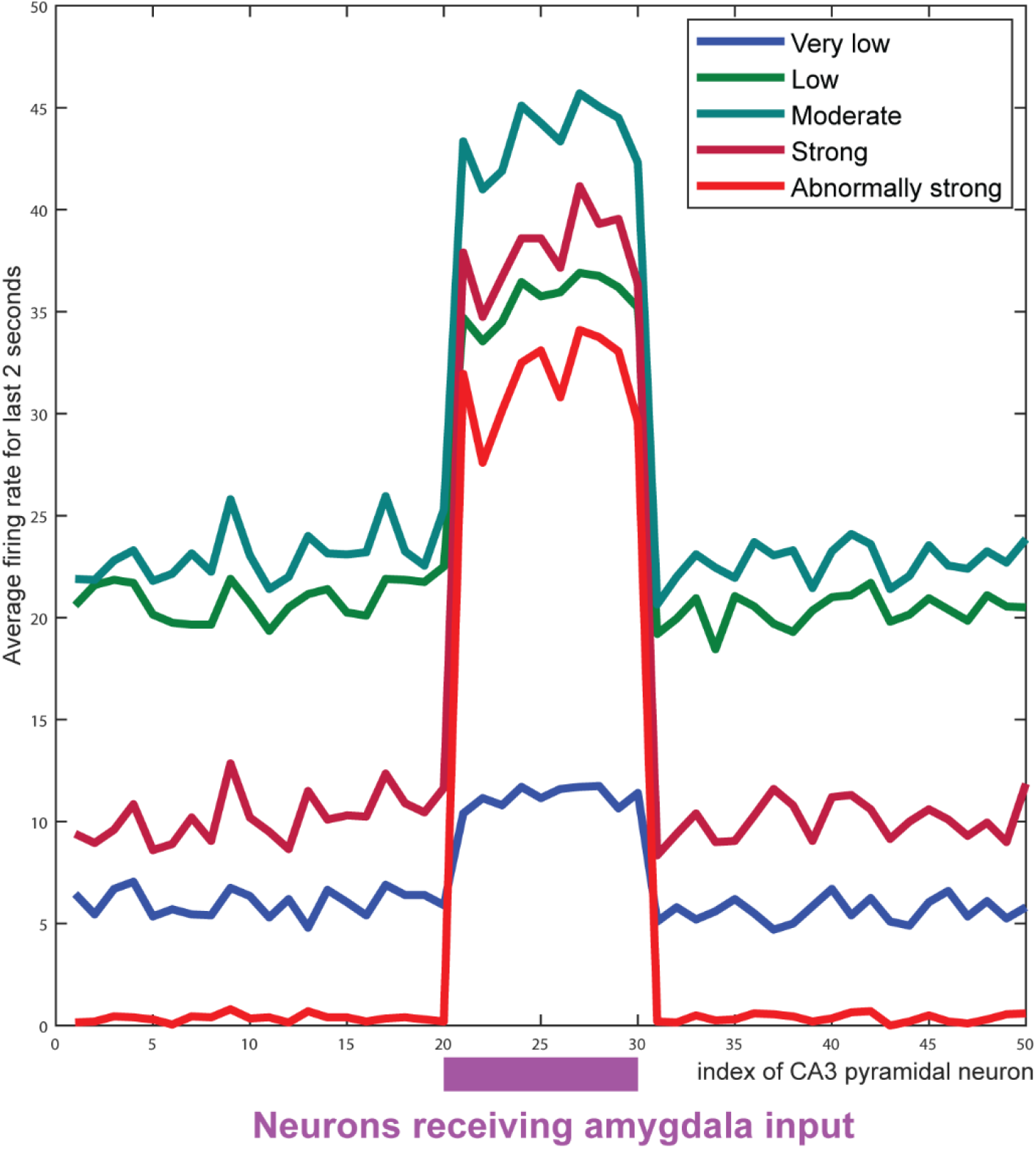
The strength of amygdalar input to CA3 affects the level of contextual detail. Each trace is computed from the average of 20 trials with different initial conditions and Poisson input trains. For each trial, the average firing rate of CA3 pyramidal neurons was computed for the last 2 seconds. For each of the five conditions shown in Figure 5, there is a corresponding trace. Neurons numbered 21 to 30 (indicated by the purple band) receive direct amygdalar input, whereas the others do not. For intermediate strengths (firing rates) of amygdalar input, the CA3 pyramidal neurons are disinhibited. When amygdalar input is very high (red trace) there is strong suppression of ensembles that do not receive direct amygdalar input (neurons 1-20 and 31-50), resulting in loss of detail.

Figure 7 summarizes the effects of the amygdala on the two CA1 pyramidal ensembles. We refer to the phasic response in CA1 as the “match mode”, as it required both priming from EC and amygdala and driving from CA3. We refer to the tonic mode as the “mismatch mode” as it arose when CA3 input did not have a matching ‘counterpart’ from the EC and the amygdala. Without amygdalar priming, the CA1 pyramidal neurons responded to CA3 inputs by entering the mismatch (tonic) mode. The CA1 pyramidal neuron activities for each ensemble (the sum of their slow synaptic output activities) were averaged over the same 20 trials used for Figure 5. In each subplot (A-E), the red trace shows the average summed activity of the CA1 pyramidal ensemble that received amygdalar input, and the blue trace shows the average summed activity of the ensemble that did not receive amygdalar input.

**Figure 7.**
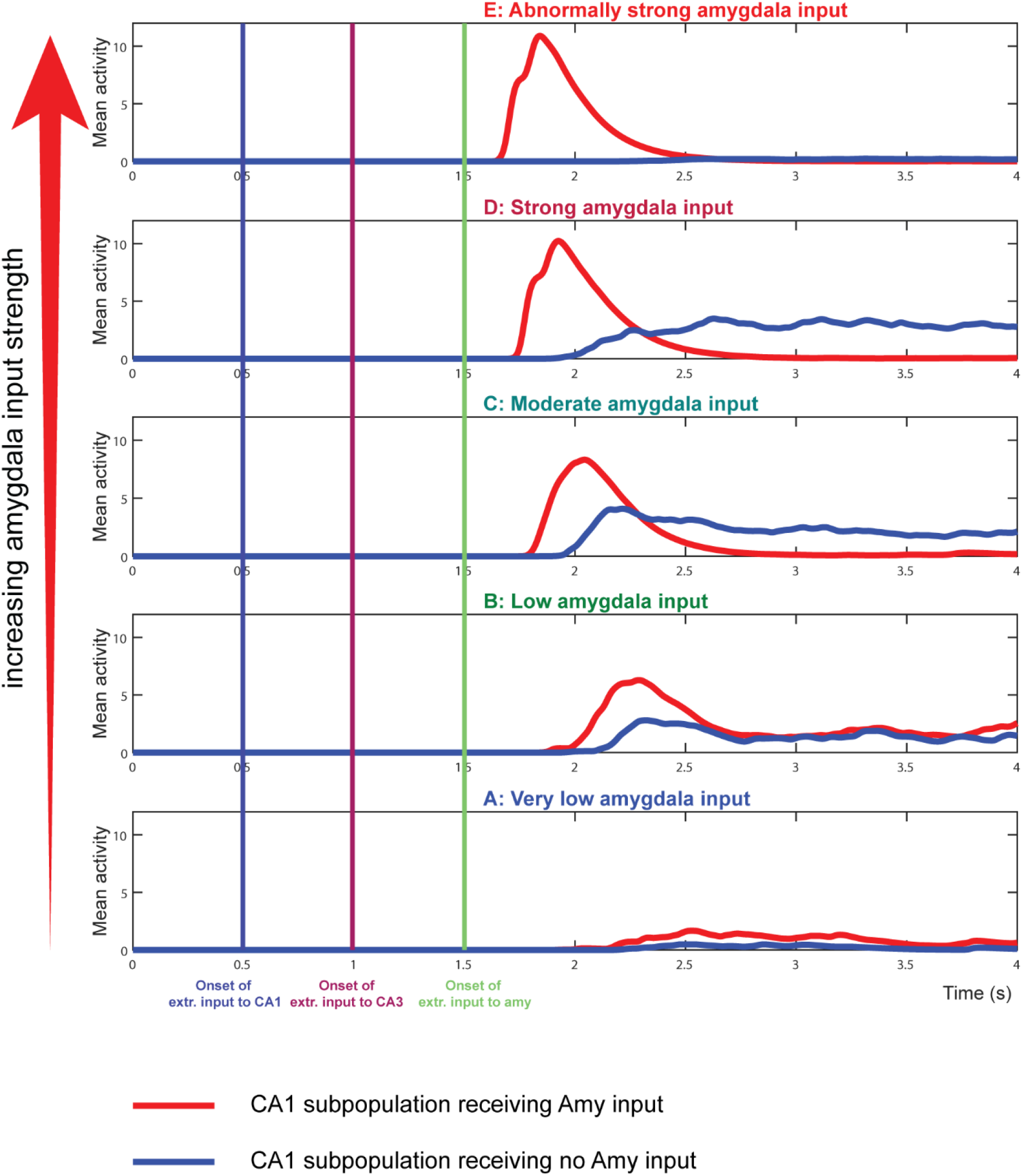
Input from the amygdala regulates the timing of CA1 output in response to a triggered memory. Each plot is computed from the average of 20 trials with different initial conditions and Poisson input trains. For each trial, the cumulative synaptic activity for the two CA1 subpopulations was computed for each of the five conditions shown in Figure 5. CA1 pyramidal neurons that receive amygdalar input respond at earlier times for stronger amygdalar input (E). Cumulative activity is computed as the sum of the slow synaptic activity generated downstream by spiking in CA1 (see Supplementary Material).

Amygdalar input also influenced the timing of hippocampal output in the match mode (Figures 5 and 7). As input strength from the amygdala increased, the phasic responses of the CA1 subpopulation that received amygdalar input occurred earlier and earlier (Figure 5; Figure 7, red traces). For the highest level of amygdalar input, the CA1 responses occurred *before* strong CA3 activity arrived in CA1, and were therefore primarily a consequence of the excessively strong priming inputs from the amygdala (Figure 5A, Figure 7A). Above a certain strength of excitation, “priming” is no longer an accurate description: it could trigger appreciable spiking in CA1 pyramidal neurons even without accompanying CA3 input.

## Discussion

Our findings reveal that amygdalar signaling to the hippocampus has three consequences. First, it enables disinhibition of memory-related patterns in CA3; second, it enables lateral inhibition of non-amygdalar input to CA3; and third, it speeds up the responses of pyramidal neurons in CA1. The first effect occurs during moderate amygdalar signaling, and corresponds to an emotion-induced enrichment of detail available for context and memory processing. The second effect occurs for abnormally strong emotion-related signaling, and masks the first effect, leading to loss of information that usually disambiguates contexts or episodes. The third effect corresponds to emotion-driven speed up in context and memory processing, which in extreme circumstances can lead to illusory categorizations of the current situation. Thus, our computational model elucidates some intriguing and testable consequences of the mechanism inferred from the distinct pattern of amygdala to hippocampus connectivity in primates (1).

Our amygdala-hippocampus circuit model performs episodic or contextual comparison, a role often attributed to the hippocampus (28,29). An active pattern in CA3 serves as a remembered template with which to compare the current situation. Our assumption is that the EC samples the current context of the organism for subsequent comparison in CA1. The CA1 pyramidal neurons either produce a phasic match signal or a tonic mismatch signal, depending on whether the CA3 driving input is accompanied by adequate EC input (24). Downstream areas can use these qualitatively distinct signals to flexibly respond to predictable versus unexpected situations. For example, a match can serve as a rapid and low-dimensional “flag” that sets off a prepared or primed response. In contrast, a mismatch may trigger a variety of responses, including alerting, attentional reorientation, or heightened learning. Such responses are likely to continue until a new match state is attained. Maintaining such responses is therefore likely to require tonic firing rather than a transient, phasic response.

In its normal operating range, the amygdala triggers disinhibition of CA3 pyramidal neurons, thereby allowing relevant memory templates to become available for comparison in CA1. This allows emotional salience to enrich hippocampal representations. The amygdala also disinhibits CA1 pyramidal neurons, facilitating their role as comparators of CA3 and EC activity. In contrast, when the amygdala activity is too strong, our simulations revealed that the effect via CA3 PV interneurons overwhelms the disinhibition via CA3 CR interneurons. This results in silencing of CA3 pyramidal neurons that do not receive direct amygdalar excitation, whose synapses are comparatively smaller (1). Direct amygdalar excitation of CA3 pyramidal neurons is very powerful, so it can overcome local inhibition from the PV interneurons. Consequently, high activation of the amygdalar pathway during intense emotional arousal activates strongly the powerful PV inhibitory interneurons, which silence nearby (non amygdalar-recipient) neurons. In this scenario, the strongly activated and powerful PV neurons act as a filter that allows only signals from the amygdala to survive in CA3 for transmission to CA1.

The latter state corresponds to an impoverished memory template – fewer features of the remembered pattern are available to CA1 for comparison with EC input. The hippocampus, in general, and CA1 in particular, has been linked with conjunctive processing of inputs (30–32). If CA1 performs comparisons of conjunctive patterns in EC and CA3, then the lower the number of features being compared, the easier it will be for matches to occur. In other words, reducing the dimensionality of CA3 mnemonic patterns can lead to overgeneralization, as more situations will trigger matches with the remembered template. If the template is a memory of a fearful or traumatic event, this can cause inappropriate generalization of fear-related memories to any stimulus that broadly and non-specifically resembles the fear-related stimulus, such as a noise.

Overgeneralization is frequently observed in the context of emotion-related disorders such as pathological anxiety and post-traumatic stress disorder (PTSD) (33–37). The hippocampus is implicated in fear-related overgeneralization, often in conjunction with the amygdala and medial prefrontal cortices (38–41). Our simulation results are in line with a recent study showing that weaker context representations prior to fear learning were predictive of later fear generalization (41). While we did not simulate learning, it is straightforward to predict how learning would be affected by an abnormally intense emotional experience, which reduces the number of active CA3 ensembles via strong inhibition from PV interneurons. If an impoverished CA3 representation serves as the basis for subsequent associative learning, overgeneralization will occur, as a result of the lack of detail in the CA3 memory pattern. This is depicted schematically in Figure 8. For example, an emotionally intense experience could lead to a failure to represent information about a situation (42,43) other than some prominent feature, such as the sound of an explosion. This condition would set the stage for subsequent association of that particular sound with fear. In support of this hypothesis, there is evidence that stress reduces the binding of temporal memories (44). The hippocampus has been associated with the learning of conjunctive representations (30–32,45–47) whereby the number of details determines its specificity. Loss of detail, which occurs in our simulations as a result of abnormally strong amygdalar input to CA3, then leads to “vague” memories. The more vague or non-specific a memory, the more frequently it can match with new circumstances. Given that amygdalar input is the strongest signal present in the low-detail state, these vague memories are likely to be of an emotional nature.

**Figure 8.**
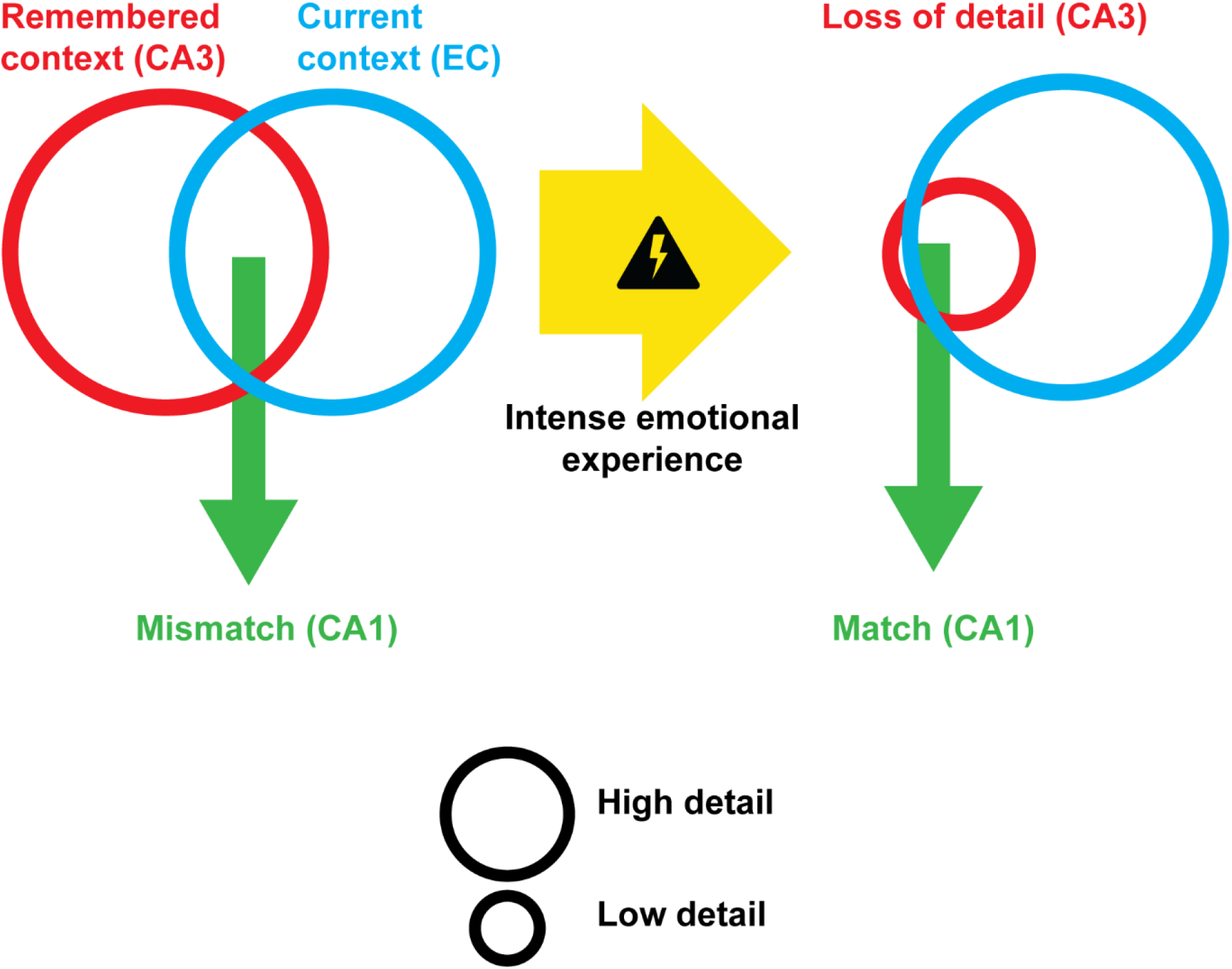
Loss of event detail in CA3 can lead to overgeneralization. When the amount of mnemonic or contextual detail is reduced in CA3 due to abnormally strong amygdalar signaling, there is an increased probability of a match computed in CA1 between the CA3 representation and the current situation, conveyed by EC. Overgeneralization has been observed in psychiatric disorders, such as phobias and PTSD.

The temporal effects of increasing amygdalar input to the hippocampus (Figure 7) may be adaptive in some circumstances. In the event of a strong emotional signal, faster categorization of the current situation may be useful, even if the learned category is less detailed. In other words, emotional salience signals conveyed to the hippocampus may speed up contextual or episodic recall and associated behavior, e.g., avoidance. But in severe cases, the ‘recall’ in CA1 actually occurs *before* any memory pattern in CA3 has been strongly activated (Figure 5E; (7E). We might think of this as a self-fulfilling prophecy in the affective domain, or as an illusory categorization.

Our study highlights how amygdalar interactions with the CA3 and CA1 subfields of the hippocampus can lead to loss of detail (in CA3) while preserving familiarity (the match response in CA1). However, this is unlikely to be the only basis for overgeneralization of context and episodic memory within the hippocampal complex. In this modeling study we have not specified the structure of the signals provided by EC to CA1 for comparison with CA3 signals, beyond assuming that it captures information about the present context of events, rather than previously learned information. Further work can explore how abnormal alterations of the signaling from EC to hippocampus may also contribute to overgeneralization, potentially via a mechanism analogous to the one demonstrated here, which is mediated by powerful synaptic influence. The present model suggests a testable mechanism for emotion-guided modulation of contextual and mnemonic information. Because disruption of the CA1 comparator leads to spurious match signals, the mismatch signals that ordinarily drive new learning are weak or absent. Thus, overgeneralizations caused by potent amygdalar inputs are also ‘sticky’, i.e., resistant to unlearning despite exposure to changed contingencies that would otherwise induce the brain to correct such overgeneralizations.

Our discussion of the implications of the model dynamics implicitly assumes that the signals from the amygdala to the hippocampus carry negative or aversive valence. However, it is known that basolateral amygdala neurons can also signal positive valence, which in turn can facilitate social interactions (48,7,6). This raises the possibility of overgeneralization caused by positive or pleasant emotions. We cannot rule out the possibility that this circuit can mediate such effects, but the importance of amygdala-hippocampus interactions in fear and anxiety is well established and clinically relevant. The framing of implications of our results in terms of fear-related overgeneralization is based on their severe clinical consequences in persistent psychiatric disorders, such as PTSD, phobias and pathological anxiety.

## Acknowledgements

The works was supported by the National Institutes of Mental Health Grant Nos. 01MH057414 and R01MH117785 [to HB].

## Disclosures

The authors reported no biomedical financial interests or potential conflicts of interest.

## Supplementary Material

### Spiking model

In both CA3 and CA1, four groups of neurons were simulated: excitatory pyramidal (Py) neurons, dendrite-inhibiting (DI) interneurons, calretinin (CR) interneurons, and parvalbumin (PV) interneurons. Each pyramidal neuron is modeled as having two compartments: one dendritic and one somatic. In addition, a population of pyramidal neurons in the amygdala (Amy) was simulated. Each neuron is modeled as a spiking Izhikevich neuron (1). The voltage in millivolts of each model neuron (or compartment) is governed by the following differential equation:

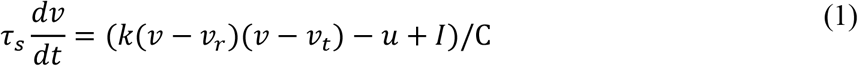

where *v* represents the voltage of the neuron or neural compartment (in millivolts), *u* is a recovery variable, and the term *I* is the total input current. The remaining terms (*k, v*_*r*_, *v*_*t*_) are parameters that facilitate approximating various types of firing pattern (Izhikevich, 2003). The recovery variable for each regular-spiking neuron (Py, DI, and CR) is governed by

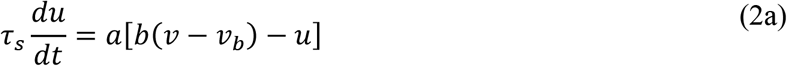

and for each fast-spiking neuron (PV) it is governed by

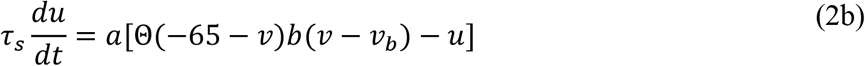

where is the Heaviside step function. Post-spike resetting is governed by the relation

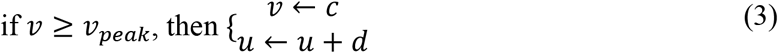

The integration time constant is τ_*s*_ (= 1 millisecond). Mean parameters for the regular-spiking neurons were a = 0.03, b = −2, c = −55, d = 100, *C* = 100, *v*_*r*=_ − 60, *v*_*t*_ = −40, *k* = 0.7, *v*_pea*k*_ = 35, *v*_*b*_ = − 60. Mean parameters for the fast-spiking neurons were a = 0.2, b = 0.025, c = −45, d = 2, *C* = 20, *v*_*r*=_ − 55, *v*_*t*_ = −40, *k* = 1.0, *v*_*peak*_ = 25, *v*_*b*_ = − 55.

### Glutamatergic and GABA-ergic currents

To model the conductances that control synaptic currents, we use the saturating differentials (SD) spike-dependent signal *gSD* (2). Thus, all synaptic conductances are governed by a couplet of differential equations of the form:

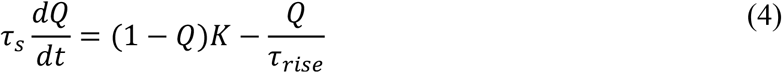

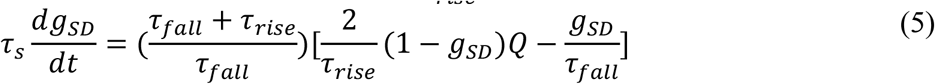

For each synapse type, an intermediate variable *Q* is triggered by the arrival of a discrete presynaptic spike *K*, which models the release of transmitter into the synaptic cleft. The spike variable *K* takes the value of 0.1 at the moment the voltage *v* goes above the spiking threshold (*v*_*peak*_), and lasts for the duration τ_*rise*_. Dynamics of different types of synapses are captured by setting values of parameters τ_*rise*_ and τ_*pfall*_, which determine the rise and fall times, respectively, of each conductance variable.

Post-synaptic currents depend on (i) summed conductance variables corresponding to each presynaptic neuron and (ii) the post-synaptic voltage. The AMPA current inputs 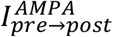 to a postynaptic neuron are computed as follows:

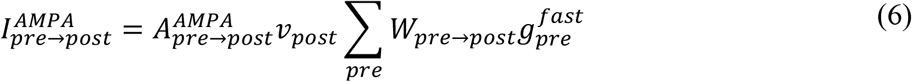

where *post* and *pre* represent the indices of the postsynaptic and presynaptic neurons, respectively, 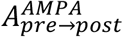 is an amplitude, and the sum is taken over weighted inputs from all presynaptic neurons. Similarly, the NMDA current inputs are computed as follows:

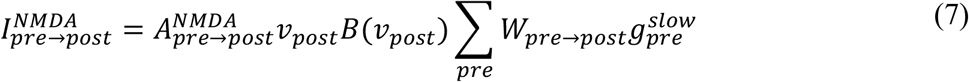

where 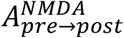 is an amplitude.

The NMDA voltage-dependence is determined by B(*v*) = [1 + 0.4202exp (0.062 ∗ *v*)]^−1^. Inhibitory GABA-ergic currents are given by the following equations:

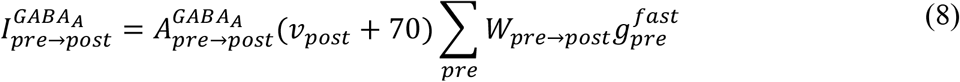

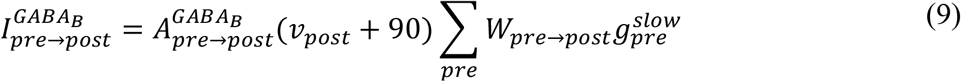

In each equation above (6-9), the adjacency matrix W_*pre*→*post*_ determines whether a connection exists. Values of W_*pre*→*post*_ are 1 or 0 depending on whether a connection exists (described below). The terms 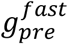 and 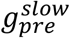 are each governed by equations (4) and (5). Each neuron therefore is associated with one fast and one slow conductance variable, which determine the effect of that neuron on downstream neurons. The rise and fall time constants (parameters τ_*rise*_ and τ_*pfall*_) for the fast conductance variables were 1 millisecond and 6 milliseconds, respectively.

For the slow conductance variables, they were 20 milliseconds and 200 milliseconds, respectively.

### Compartmental coupling

Electrical coupling between two compartments (indexed 1 and 2) is given by

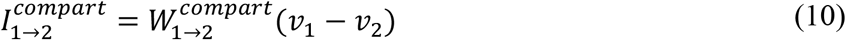

where the subscripts 1 and 2 represent the compartment index, which corresponds to either soma or dendrite. Pyramidal cells in CA3 and CA1 each possess these 2 compartments. Dendrites are modeled as active compartments, using the regular-spiking parameters.

### KCNQ channel

The CA1 Py neurons also possess KCNQ channels, which generate the M-current that enables match/mismatch responses. Our M-current is derived from the detailed biophysical model of Berteau and Bullock (3). The current is given by

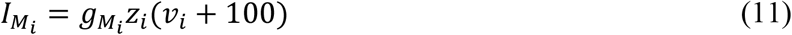

where *v*_*pi*_ is the voltage of CA1 Py neuron, 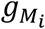 is the maximal conductance of the KCNQ channel and *z*_*i*_ is the gating variable, governed by

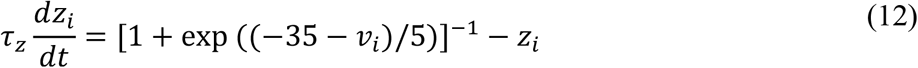

where time constant τ_*pz*_ = 200 milliseconds. The 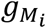 value is higher for the somatic than the dendritic compartment.

### Connectivity

The CA3 subfield contains 50 Py, 20 PV, 20 DI, and 20 CR neurons. The CA1 subfield contains 100 Py, 20 PV, 20 DI, and 20 CR neurons. In addition, 10 Amy neurons were simulated.

Baseline inputs to each neural group, as well as input from the entorhinal cortex (EC), were simulated as Poisson spiking input. The CA3 subfield is divided into five ensembles containing equal numbers of each type of neuron. Similarly, the CA1 subfield is divided into two subgroups containing equal numbers of each type of neuron.

In CA3, only the 3^rd^ ensemble received Amy input. Py neurons excited PV interneurons within their own ensemble. DI interneurons inhibited Py neurons within their own ensemble. CR interneurons inhibited DI interneurons in all ensembles, enabling disinhibition of all Py neurons. PV interneurons inhibited Py neurons in all ensembles, enabling both lateral or “off-surround” inhibition and feedback or “off-center” inhibition. PV interneurons also inhibited each other.

In CA1, Amy input was received by all CR interneurons, disinhibiting priming input from EC to all Py neurons. Only the first ensemble of CA1 Py neurons received Amy input. PV interneurons inhibited Py neurons and other PV interneurons in both ensembles. CA3 Py neurons sent excitation to all CA1 Py neurons and PV interneurons. Py neurons excited PV interneurons within their own ensemble.

For each excitatory connection, the AMPA:NMDA ratio 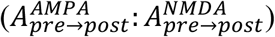 was set to 1 :

0.3; for each inhibitory connection, the GABA: GABA ratio 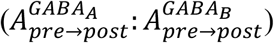 was set to 1 : 0.1. Values of terms in the adjacency matrices W_*pre*→*post*_ are 1 or 0 depending on whether a connection exists. The strength of an existing synaptic connection is given by the parameters A_*pre*→*post*_ in Table 1.

**Table 1.** Parameters used in the simulations. Each trial consisted of 5 simulation runs. For a given trial the same connection weight matrices were used in all simulations. Connection weights were drawn from a uniform distribution with the mean value as specified in the table, and bounded between ±2.5% of the mean.

| Parameter | Mean Value | Parameter | Mean Value |
| --- | --- | --- | --- |
| $A_{input \rightarrow Amy}^{AMPA}$ | 50 | $A_{CA3DI \rightarrow CA3PyD}^{GABA_A}$ | 60 |
| $A_{input \rightarrow CA3PyD}^{AMPA}$ | 50 | $A_{CA3CR \rightarrow CA3DI}^{GABA_A}$ | 40 |
| $A_{input \rightarrow CA3DI}^{AMPA}$ | 50 | $A_{CA3PV \rightarrow CA3PV}^{GABA_A}$ | 5 |
| $A_{input \rightarrow CA3PV}^{AMPA}$ | 20 | $A_{CA3PV \rightarrow CA3PyS}^{GABA_A}$ | 11 |
| $A_{input \rightarrow CA3CR}^{AMPA}$ | 30 | $A_{CA1D \rightarrow CA3PyD}^{GABA_A}$ | 15 |
| $A_{input \rightarrow CA1PyD}^{AMPA}$ | 6 | $A_{CA1CR \rightarrow CA1DI}^{GABA_A}$ | 4 |
| $A_{input \rightarrow CA1DI}^{AMPA}$ | 20 | $A_{CA1PV \rightarrow CA1PyS}^{GABA_A}$ | 0.9 |
| $A_{input \rightarrow CA1PV}^{AMPA}$ | 10 | $A_{CA1PV \rightarrow CA1PV}^{GABA_A}$ | 0.25 |
| $A_{input \rightarrow CA1CR}^{AMPA}$ | 20 | $g_{M_{CA1PyD}}$ | 2 |
| $A_{CA3PyS \rightarrow CA1PyS}^{AMPA}$ | 6 | $g_{M_{CA1PyS}}$ | 40 |
| $A_{CA3PyS \rightarrow CA3PV}^{AMPA}$ | 0.25 | $W_{PyS \rightarrow PyD}^{compart}$ | 9 |
| $A_{Amy \rightarrow CA3PyD}^{AMPA}$ | 20 | $W_{PyD \rightarrow PyS}^{compart}$ | 21 |
| $A_{Amy \rightarrow CA3CR}^{AMPA}$ | 6 | $P_{CA3PyD}$ | 200 |
| $A_{Amy \rightarrow CA3PV}^{AMPA}$ | 6 | $P_{CA3DI}$ | 400 |
| $A_{Amy \rightarrow CA1PyD}^{AMPA}$ | 4 | $P_{CA3CR}$ | 10 |
| $A_{Amy \rightarrow CA1CR}^{AMPA}$ | 10 | $P_{CA3PV}$ | 20 |
| $A_{CA1PyS \rightarrow CA1PV}^{GABA_A}$ | 0.075 | $P_{CA1PyD}$ | 150 |
| | | $P_{CA1DI}$ | 60 |
| | | $P_{CA1CR}$ | 4 |
| | | $P_{CA1PV}$ | 20 |

### Input timing

Extrinsic inputs to each neuron group take the form of Poisson spike trains. Here “extrinsic” refers to excitatory inputs originating in neurons that have not been explicitly modeled as Izhikevich neurons. These inputs include the non-amygdala inputs to CA3 Py neurons and interneurons, the EC inputs to CA1, and the inputs to the Amy neurons. As mentioned in the main text, the onset times of these extrinsic inputs are staggered. EC input to CA1 is turned on at 0.5s, input to CA3 at 1s, and input to Amy at 1.5s. The Poisson rate of input to the Amy neurons increases from the low input simulations (Figure 5A) to the abnormally strong input simulations (Figure 5E). All other Poisson rates are constant for every simulation. The excitatory inputs to the CA3 Py neurons are unable to trigger firing until disinhibition occurs via CR interneurons, which are excited by Amy neurons. The connection weights are set up so that for moderate Amy input and higher, the DI interneurons are completely silenced by the CR interneurons, and thus the CA3 Py neurons receive their maximal excitatory input. Further increases in Amy activity recruit PV interneurons, which cause perisomatic inhibition. Given that each CA1 Py neuron receives driving input from all the CA3 Py neurons, the timing of onset of CA1 Py neurons is modulated by the total strength of CA3 Py neurons. Poisson input rates *P* in spikes per second are specified in Table 1.

The order of onset times of extrinsic excitatory inputs was chosen based on considerations of behavior and hippocampal connectivity. The excitatory inputs to the CA3 are assumed to arrive after extrinsic excitatory inputs to CA1. Our assumption is that the majority of CA3 activation occurs through a disynaptic pathway via EC and dentate gyrus, whereas CA1 activation occurs through a monosynaptic pathway from EC. Thus, if a common EC signal is the source of both CA3 input and CA1 input, the priming inputs to the dendritic compartment of CA1 Py neurons would be expected to arrive before the inputs that drive CA3 Py neurons. This ensures that the driving inputs to the somatic compartment of CA1 Py neurons occurs after the priming input. If the priming input to CA1 is sufficiently early, the match mode becomes possible.

### Averaging for summary plots of CA1 (Figure 7)

For every spike generated in a model Py neuron in CA1, there was a resultant slow synaptic output signal g_i_^*slow*^ received by the Py neuron’s targets (e.g., CA1 PV interneurons). A measure of the total activity of the two CA1 subpopulations is computed by summing these slow synaptic output signals for all Py neurons in a subpopulation within a given simulation. Each simulation corresponds to a level of amygdala input to the circuit. The average of this sum over all trials was computed for each simulation.

The single-simulation total activity at each time point 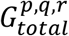 is given by

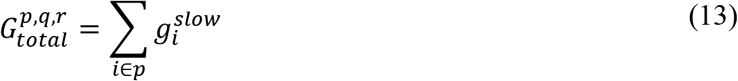

where *p* refers to the specific CA1 subpopulation, which is either the one that receives input from Amy, or the one that does not. The variable 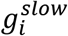 is governed by equations (4) and (5), and determines the slow component of a CA1 neuron’s impact on downstream neurons. It functions effectively as a smoothed version of the spike rate of the corresponding CA1 neuron. Superscript *q* refers to the simulation number, ranging from 1 to 5, in which the strength of input to amygdala is varied. Superscript *r* refers to the trial number.

The multi-trial average activity of each subpopulation at each time point is then given by

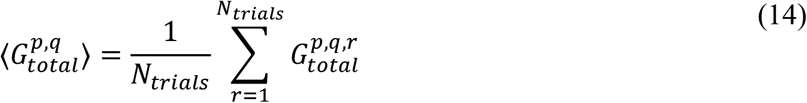

where *N*_*trials*_(= 20) is the total number of trials. This formula determines the red and blue traces shown in Figure 7. The red traces correspond to the subpopulation that receives input from amygdala, and the blue traces correspond to the subpopulation that does not.

### Simulation details

To establish the robustness of the qualitative effects, we tested 20 trials with different initial conditions. In each trial, 5 simulations were run with different levels of input to Amy neurons (Poisson rates *p*: 12.5, 25, 50, 100 and 200 spikes per second). The Izhikevich parameters, synaptic time constants and connection weights were each drawn from a random distribution centered on the mean value specified above, and bounded between ±2.5% of the mean.

Simulations were performed in MATLAB R2021b. Differential equations were solved using the forward Euler method with a time step of 0.25 milliseconds. Each simulation corresponded to 8 seconds of real time.

## Notes

### Competing Interest Statement

The authors have declared no competing interest.

